# Laboratory and field trials reveal the potential of a gel formulation of entomopathogenic nematodes as biocontrol against the fall armyworm (*Spodoptera frugiperda*)

**DOI:** 10.1101/2022.02.03.479057

**Authors:** Patrick Fallet, Didace Bazagwira, Julie Guenat, Carlos Bustos-Segura, Patrick Karangwa, Ishimwe Primitive Mukundwa, Joelle Kajuga, Thomas Degen, Stefan Toepfer, Ted C.J. Turlings

## Abstract

1.

The fall armyworm (FAW), *Spodoptera frugiperda* (Lepidoptera: Noctuidae) can cause tremendous yield losses in maize. Its invasion into Africa and Asia has led to an enormous influx of insecticides into maize agro-ecosystems. Safe, effective and readily available alternatives are desperately needed. Entomopathogenic nematodes (EPN) are commonly used against soil insect pests, but can also control some above-ground pests. We explored the possibility to control FAW by incorporating EPN into a formulation that can be easily applied into the whorl of maize plants, where the caterpillars mostly feed. We tested this approach in laboratory cage experiments as well as in field trials. In the laboratory, treating maize plants with a low dose of EPN in a carboxymethyl cellulose (CMC) gel formulation (about 3000 infective juveniles per plant), caused 100% mortality of FAW caterpillars and prevented plant damage considerably, whereas EPN applied in water or a surfactant-polymer-formulation (SPF) caused 72% and 94% mortality, respectively. Under field conditions, one-time treatments with EPN applied in water, SPF or CMC gel were all able to prevent significant plant damage, but only the EPN-gel formulation significantly reduced FAW infestation. Notably, the gel formulation was as effective as a standard dose of cypermethrin, an insecticide commonly used against FAW. Repeated applications may be needed to reduce re-infestations by FAW across a whole cropping season depending on the local maize phenology and pest dynamics. These findings demonstrate that EPN are excellent candidates for the biological control of FAW and are a safe and sustainable alternative to chemical insecticides.

**Highlights:**

- Entomopathogenic nematodes are highly lethal to fall armyworm caterpillars.
- Appropriate formulation of the nematodes is crucial for above-ground application.
- A gel formulation of entomopathogenic nematodes was as effective as chemical insecticides.
- Entomopathogenic nematodes can be used for the control of fall armyworm in maize.

## 2. Introduction

The fall armyworm (FAW), *Spodoptera frugiperda* (Lepidoptera: Noctuidae) is native to the tropical and subtropical regions of the Americas (Luginbill, 1928). Although FAW is polyphagous and can feed on more than 350 plant species (Montezano et al., 2018), it has a clear preference for maize, sorghum, rice and some other grasses (Luginbill, 1928). In early 2016, FAW was observed for the first time in western Africa (Goergen et al., 2016) and soon after spread throughout much of sub-Saharan Africa (Cock et al., 2017; Day et al., 2017). In mid-2018, FAW was first discovered in India and rapidly spread throughout southern and eastern Asia (Sharanabasappa et al., 2018). FAW is predicted to spread even further (Early et al., 2018; Liu et al., 2020).

FAW frequently causes severe damage to maize and can substantially reduce yields, causing tremendous economic losses (Baudron et al., 2019; Day et al., 2017; Hruska and Gould, 1997; Rwomushana et al., 2018; Wan et al., 2021). Due to its voracious and fast feeding behaviour as well as its exceptional migration capabilities, it currently threatens the food security of millions of people (Babendreier et al., 2020; Day et al., 2017; Rwomushana et al., 2018). To mitigate the impact of FAW, several control options are applied including synthetic insecticides, genetically modified crops that contain *Bacillus thuringiensis* (*Bt*) toxins, biopesticides (e.g. virus, fungus, bacteria or nematodes), botanicals (e.g. neem extract), mechanical control (e.g. handpicking caterpillars), or special agronomic practices (e.g. push and pull) (Abrahams et al., 2017; Guo et al., 2020; Harrison et al., 2019; Wan et al., 2021). However, chemical insecticides have quickly become the backbone of FAW control in Africa and Asia, mainly due to unavailability of alternatives and due to governmental emergency programmes subsidising synthetic insecticides (Abrahams et al., 2017; Tambo et al., 2020). This situation has led to an enormous influx of insecticides in previously rarely treated maize-growing areas (Yang et al., 2021). High frequency and broad-scale use of synthetic insecticides can have negative consequences for human health and the environment. It also substantially reduces populations of beneficial natural enemies and leads to resistance in populations of FAW, an obvious disadvantage compared to biological control agents. FAW resistance had already been reported to a variety of chemical insecticides (Wan et al., 2021), as well as to single *Bt* toxic proteins (Blanco et al., 2016; Farias et al., 2014; Storer et al., 2010). Hence, there is an urgent need for readily available, safe, effective, and sustainable alternatives (Day et al., 2017).

Entomopathogenic nematodes (EPN) are tiny soil dwelling roundworms that can be found naturally in soils worldwide (Hominick et al., 1996). EPN can infest and kill a large variety of insects (Kaya and Gaugler, 1993). Many EPN species or strains are highly virulent to lepidopteran larvae, including FAW (Acharya et al., 2020; Andaló et al., 2010; Caccia et al., 2014; Fuxa et al., 1988; Kaya and Gaugler, 1993; Richter and Fuxa, 1990). Unlike many pesticides, EPN pose no risk to farmers or consumers, and hardly any risk to the environment (Ehlers and Hokkanen, 1996). They can be mass-produced (Ehlers, 2001) – also in Africa (Holmes et al., 2015) – and have the potential to be cost effective if formulated and applied correctly (Ehlers, 2001; Kagimu et al., 2017).

Due to limited survival of EPN outside the soil, they have mainly been used against soil pests (Kagimu et al., 2017; Kaya and Gaugler, 1993; Lacey and Georgis, 2012). As soil-dwelling organisms, EPN are highly susceptible to desiccation, ultraviolet radiation and heat (Kagimu et al., 2017; Kaya and Gaugler, 1993; Lacey and Georgis, 2012), which limits their use against aboveground pests. To resolve these limitations, EPN have been, as many other biopesticides and chemicals, incorporated into formulations that protect them against these abiotic factors (Beck et al., 2013; Glazer et al., 1992; Glazer and Navon, 1990; Hiltpold, 2015; Navon et al., 1998; Schroer et al., 2005; Shapiro-Ilan et al., 2010; Shapiro-Ilan et al., 2012). Each formulation has its advantages and disadvantages, and each plant-pest system may need its own optimized solution.

We hypothesised that the maize–FAW pest system may be particularly well-suited for the application of EPN because FAW caterpillars mostly feed in protected areas such as deep in the wrapped leaves of the whorl or on the cob under the husk leaves (Buntin, 1986; Labatte, 1993; Luginbill, 1928). Although well-suited for EPNs, such feeding behaviour makes FAW caterpillar control difficult with conventional flat sprays of pesticides, particularly when direct contact is required (Pannuti et al., 2015). In contrast, EPN applied into the whorl of maize or directly onto the cobs will be able to actively forage for FAW caterpillars. Moreover, the leaves will protect the EPN from unfavourable abiotic factors, providing higher humidity, reduced temperature and less radiation exposure, as compared to an open surface. In order to ensure that EPN are even better protected as well as to enhance their longevity and finally to assure good control efficacy, we aimed at incorporating EPN into formulations that are particular suitable for application onto maize plants. In a preliminary study, we formulated EPN in sand as well as in two types of alginate beads, but after unsatisfactory results, switched to a more detailed study using a commercial surfactant-polymer-formulation and a carboxymethyl cellulose-based gel. We tested those formulations in several series of laboratory experiments and evaluated the most promising ones in field trials. The here reported findings should significantly advance the development of a formulation that will offer practitioners a way to achieve safe, sustainable and effective control of FAW using EPN.

## 3. Material and methods

### 3.1. Origin and handling of nematodes

Two entomopathogenic nematode species were used in this study. These were *Steinernema carpocapsae* (strain RW14-G-R3a-2) and *Heterorhabditis ruandica* (strain Rw18_M-Hr1a;), both being among the strains isolated from soil samples in Rwanda in 2014 and 2018, respectively (Fallet et al., 2020; Machado et al., 2021; Yan et al., 2016). They were among the most effective strains in killing FAW caterpillars in a screening test where we compared 40 EPN strains, representing twelve species, originating from Rwanda, Mexico and from a few commercial sources (Fallet et al., in press). EPN were reared *in vivo* on larvae of *Galleria mellonella* L. (Lepidoptera: Pyralidae) and stored in darkness at 12°C. They were used within a week post emergence from *G. mellonella* cadavers.

### 3.2. Tested formulations

In preliminary laboratory assays (Fig. S2-S5), we compared the efficacy of eight EPN-formulations in killing FAW caterpillars on potted maize seedlings. The formulations tested were: (1) water, (2) sand, (3) alginate beads (as described in Kim et al., 2021), (4) a commercial bead (Nema-Caps®, Agrocaps SPRL, Gedienne, Belgium), (5) Navon’s alginate gel (as described inNavon et al., 2002), (6) a gel made from carboxymethyl cellulose (CMC) (Sigma-Aldrich, St. Louis, MO, USA), and two commercial liquid formulations, (7) an emulsifiable vegetable oil (Addit^®^, Koppert Biological Systems, Berkel en Rodenrijs, Netherlands) and (8) a surfactant-polymer-formulation (SPF) (Nemaperfect^®^, e-nema GmbH, Schwentinental, Germany). Based on these preliminary trials, we concluded that the most promising formulations were water (for its low cost and ease of use), the CMC gel (for its high efficacy) and the commercially available SPF (Nemaperfect®, for its ease of use and efficacy). These three formulations were further investigated in this study.

SPF and CMC were dissolved in water to final concentrations of 0.2 % and 3 % respectively by rapid stirring in a 200 mL beaker until complete dissolution. The two formulations were first completely dissolved in 90 mL of water, then 10 mL of concentrated freshly emerged infective juveniles (IJs, the free-living stage of EPN; < 1 week old, 150’000 EPN) were added. The same procedure was used to incorporate EPN in just water. Using a stereoscopic microscope, we confirmed that the formulations contained approximately 1500 IJs/mL. Formulations were kept in cool boxes until use, which occurred within 30 minutes in laboratory experiments or up to a few hours in the field experiments.

### 3.1. Efficacy of EPN formulations in reducing fall armyworm infestation and preventing maize damage under laboratory conditions

#### 3.1.1. Maize and fall armyworm

Maize plants (Hybrid CML203 x CML204, Rwanda Animal and Agricultural Resource Board, Huye, Rwanda) were grown in plastic pots (12 cm diameter x 8.5 cm height) using commercial potting soil (Classic, Einheitserdewerke Patzer, Sinntal-Altengronau, Germany) between March 6^th^ and March 27^th^ 2020. Plants were grown for four weeks in a greenhouse supplemented with artificial light (16:8 h L:D, approx. 350 μmol * m^-2^ * sec^-1^). They were watered twice a week with water supplemented with fertilizer as specified by the supplier (engrais liquid universel, Capito, Intercoop House & Garden Cooperative, Biel, Switzerland), until they carried three to four fully developed true leaves (ca. 30-35 cm in height).

*Spodoptera frugiperda* caterpillars (FAW) were obtained from a colony at the University of Neuchâtel reared on artificial diet (Beet Armyworm Diet, Frontier Scientific, Newark, USA) under quarantine conditions (OFEV permit A140502).

#### 3.1.2. Experimental procedure

The efficacy of *S. carpocapsae* (strain RW14-G-R3a-2) against FAW was evaluated in cage experiments under laboratory conditions. Two maize plants (Rwandan hybrid CML203 x CML204, Rwanda Animal and Agricultural Resource Board, Huye, Rwanda; 30-35 cm in height) were placed inside a net cage (60 cm in height x 40 cm in depth x 40cm in width, Fig. 1) supplemented with artificial LED light (16:8 h L:D, approx. 300 μmol * m^-2^ * sec^-1^). Three third-instar FAW caterpillars (ca. 1cm in length) were placed into the whorl of each plant. Twenty-four hours later, we applied 2 mL of a given formulation into the whorl of the two plants in a cage. EPN treatments consisted of ~3000 IJs applied in 2 mL of either water, 0.2 % SPF or 3% CMC gel. As controls, we treated plants with the same three formulations but without EPN. Every morning, 2 mL of water was vaporized above the whorl (ca. 15 cm distance) to mimic the effect of the dew. Six days post treatment, we evaluated plant damage using the Davis scale (Davis whorl & furl damage scale; Davis et al., 1992) as described in Toepfer et al. (2021), where a score of “0” represent an intact plant while a score of “9” represent a plant that is almost totally destroyed. Subsequently we counted the number of surviving caterpillars. Five cages (ten plants) per treatment were used in each of three independent experiments (n = 15 cages; 30 plants per treatment).

**Figure 1:**
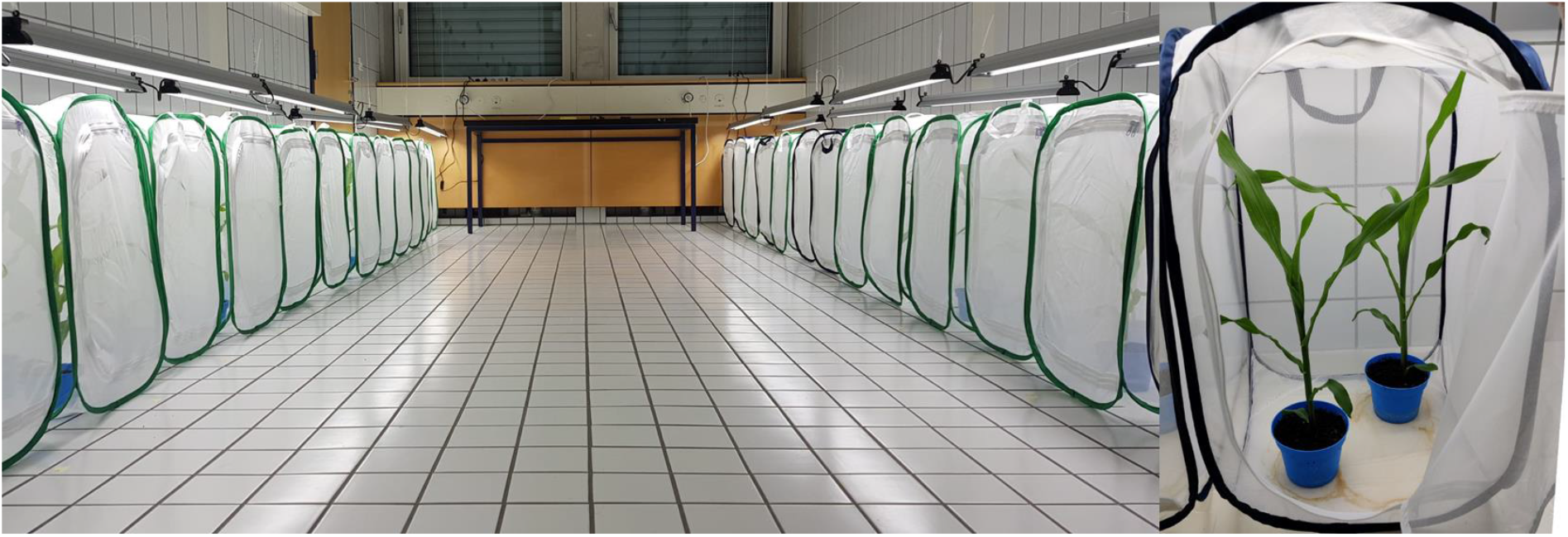
Laboratory experimental setup. Two plants were placed in a net cage and subsequently infested by placing three third-instar caterpillars into the whorl. Twenty-four hours later, a given formulation was applied into the whorl. Five cages (ten plants) were used per treatment in each of three independent experiments.

#### 3.1.3. Data analyses

Statistical analyses were performed using R version 4.1.2 (R Core Team, 2021). Given the complete separation of the data (100 % mortality in the gel+EPN treatment), we analysed caterpillar survival per cage using a General Independence Test (“coin” package; Hothorn et al., 2006). The survival of caterpillars in a cage was used as the response variable, while treatment was used as a fixed factor. Pairwise Two-Sample Permutation Tests (“rcompanion” package; Mangiafico, 2021) were used to compare treatments and were corrected for false discovery using the Benjamini & Hochberg method (1995). The efficacy of each formulation in killing FAW was estimated (Eq. B.1) as compared to their respective negative control (formulation without nematodes).

The effect of treatments on plant damage was analysed using cumulative link mixed models (“ordinal” package; Christensen, 2019), followed by multiple comparisons (“emmeans” package; Lenth, 2021) corrected for false discovery using the Benjamini & Hochberg method (1995). The damage score given to a plant was used as the response variable, while treatment was used as a fixed factor and cage as a random factor (two plants per cage). To compare the proportion of plants with medium and high damage (Davis score higher than “3”) among treatments, we used a General Independence Test (Hothorn et al., 2006)(“coin” package; Hothorn et al., 2006) and Pairwise Two-Sample Permutation Tests (“rcompanion” package; Mangiafico, 2021) corrected for false discovery using the Benjamini & Hochberg method (1995). The proportion of damaged plants was used as the response variable, while treatment was used as a fixed factor. The efficacy of each formulation in preventing medium to heavy damage as compared to their respective control (formulation without nematodes) was estimated (Eq. B.2).

### 3.2. Efficacy of EPN formulations in reducing fall armyworm infestation and preventing plant damage under field conditions

#### 3.2.1. Field sites

We assessed the efficacy of formulations containing EPN in four maize fields in Southern Rwanda. Two fields (I and II) were located at the RAB Rubona Station in the district of Huye (GPS: S 02°28.827’, E 029°45.825’; altitude 1660 m.a.s.l.) and the other two (III and IV) in the district of Nyamagabe (GPS: S 02°28.539’, E 029°28.515’; altitude 2000 m.a.s.l.). During our experiment from mid-February to early-March 2020, the mean temperature recorded at the Rubona site was 21.3±6.6°C (mean ± sd; max = 38.5°C; min = 13.3°C) and was 20.3±7.3°C (mean±sd; max = 38.7°C; min = 7.3°C) at the Nyamagabe site. At the Rubona site, the mean daily rainfall was 2.5±6.2mm (mean±sd; max = 22.1 mm; min = 0 mm). Rainfall could not be measured at the Nyamagabe site.

#### 3.2.2. Maize and fall armyworm

We planted maize (Rwandan hybrid CML203 x CML204, Rwanda Animal and Agricultural Resource Board, Huye, Rwanda) in the four fields measuring approximately 25 m by 25 between the 9^th^ and 22^nd^ of January 2020. Each field was fertilized with about 300 kg of manure. Maize plants were sown every 30 cm in rows separated by 70 cm, representing about 47’000 plants per hectare. Plants were grown for four to five weeks and were not treated with pesticides to ensure natural infestation by FAW. At the start of the experiment (before treatment), the maize plants in field I, II, III and IV measured on average 28±6.7, 23±7.7, 13±3.1 and 33±9.9 cm in height (mean±sd) and carried on average 7.8±1.6, 7.7±1, 5.9±0.7 and 9.1±1.1 leaves (mean±sd), respectively. They were found to be infested by 1.3±1.1, 1.5±1.4, 0.15±0.4, 1±0.99 FAW caterpillars per plant (mean±sd) in field I, II, III and IV, respectively, not considering neonates.

#### 3.2.3. Experimental procedure

We assessed the efficacy of formulations containing EPN in the four fields using a block design (Fig. 2). To account for varying environmental conditions (i.e. surrounding habitats, exposition, etc...) as well as FAW infestation densities across the fields, each field was divided into four quadrants (see data analysis below). The quadrants were subdivided into six plots (3.3 m x 2.1 m) of each 48 plants. All plants within a plot were treated with one of six treatments (n = 4 plots per treatment per field; 16 plots per treatment in total). Plots were separated from each other by two untreated rows of plants that served as buffer (Fig. 2). Treatments were applied as 2 mL spot spray into the whorl. Treatments consisted of: *S. carpocapsae* RW14-G-R3a-2 formulated in either water (1), in 0.2 % SPF (2), or in 3% CMC gel (3), *H. ruandica* Rw18_M-Hr1a in water (4), 5% cypermethrin (5) as positive control and water without nematodes (6) as negative control. The EPN formulations were prepared and applied as described above and contained 1’500 IJs/mL. The pyrethroid insecticide cypermethrin (Supra EC 50 g a.i. / litre, thus ca 5% a.i. in product, ETG inputs Ltd, India) was dissolved in water to a solution of 1.875 μL/mL (0.19 μg active ingredient per plant in a 2 mL spot spray).

**Figure 2:**
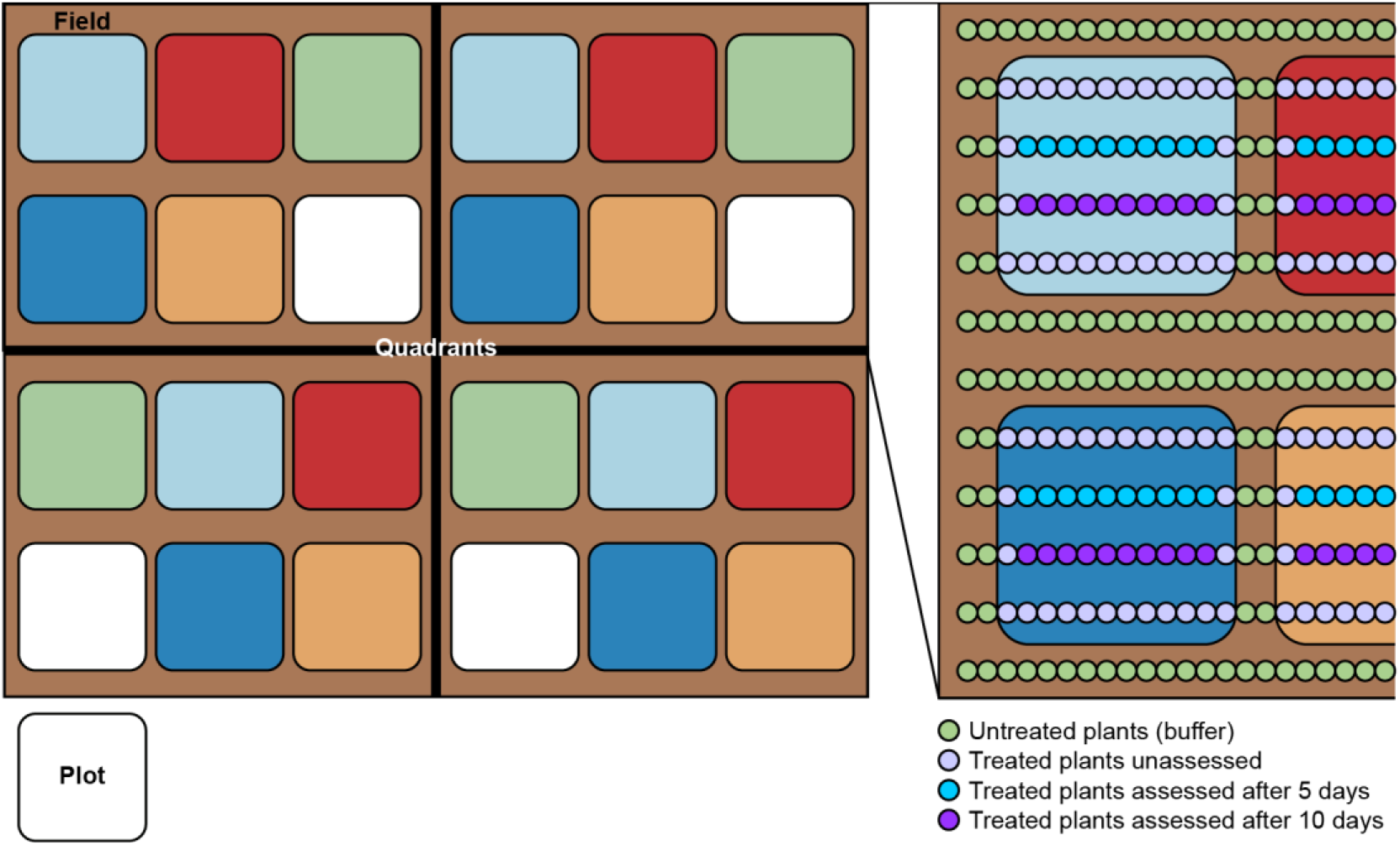
Experimental setup. Each field measured 25 m by 25 m. Maize was sown in 24 rows separated by 70 cm. Within rows, plants were separated by 30 cm. The fields were divided into four quadrants to account for the spatial variation of FAW across the fields. Quadrants were subdivided into six plots which were treated with a different treatment (coloured squares). Four plots were used per treatment in each field (squares of the same colour). The 48 plants in a plot were treated, but only ten plants inside one of the two inner rows of each plot were evaluated for damage and infestation by FAW at five (turquoise dots) and ten (purple dots) days post treatment. Two untreated rows of plants were kept between plots (green dots) as buffer.

We evaluated treatment efficacy in ten plants from one of the inner rows of the plot at five and ten days post treatment, as indicated in Fig. 2. Plant damage was assessed using the Davis scale (Davis whorl & furl damage scale; Davis et al., 1992) as described in Toepfer et al. (2021). Then we searched for surviving caterpillars. The occurrence (presence or absence) of small caterpillars (< 0.5 cm) in each plant was recorded as a proxy for re-infestation (after treatments) by FAW, while the number of older caterpillars (caterpillars longer than 0.5 cm) was used to determine remaining FAW infestation levels as a proxy for treatment efficacy.

#### 3.2.4. Data analyses

Statistical analyses were performed using R 4.1.2 (R Core Team, 2021). The number of FAW caterpillars per plot were analysed using generalized linear mixed-effects models (“lme4” package; Bates et al., 2015) with a negative binomial error distribution, followed by multiple comparisons corrected for false discovery using the Benjamini & Hochberg method (1995). The number of caterpillars per plot was the response variable. Treatment and field number were used as fixed factors, while quadrant was used as a random factor to account for the variation in the spatial distribution of FAW across the fields.. The efficacy of each treatment in reducing FAW infestation as compared to the water control (water without nematodes) was estimated (Eq. B.3).

Plant damage was analysed using cumulative link mixed models (“ordinal” package; Christensen, 2019), followed by multiple comparisons (“emmeans” package; Lenth, 2021) corrected for false discovery using the Benjamini & Hochberg method (1995). The proportion of plants with a damage score higher than “3” as well as the re-infestation by neonates were analysed using generalized linear mixed-effects models (“lme4” package; Bates et al., 2015) with a binomial distribution, followed by multiple comparisons (“emmeans” package; Lenth, 2021) corrected for false discovery using the Benjamini & Hochberg method (1995). In these tests, the response variable was either the damage score attributed to each plant or the presence/absence of neonates on each plant. Treatment and field number were used as fixed factors, while quadrant and plot were used as random factors.

## 4. Results

### 4.1. Efficacy of EPN formulations in reducing fall armyworm infestation and preventing maize damage under laboratory conditions

Treating maize with formulated-EPN significantly reduced the number of surviving FAW caterpillars under controlled laboratory conditions (*MaxT* = 4.7, *p* < 0.001; Fig. 3). The best result was obtained with 3000 IJs formulated in the CMC gel (0 % survival [mean]; Gel+EPN vs SPF+EPN: *p* = 0.23; Gel+EPN vs Water+EPN: *p* = 0.002; Gel+EPN vs Gel: *p* < 0.001), closely followed by the commercial SPF (4±13 % survival [mean±sd]; SPF+EPN vs Water+EPN: *p* = 0.02; SPF+EPN vs SPF: *p* < 0.001). EPN applied in just water was the least effective of the EPN formulations (21±20 % survival [mean±sd]; Water+EPN vs Water: *p* < 0.001). The formulations without EPN did not affect fall armyworm survival (Water vs SPF: *p* = 0.7; Water vs Gel: *p* = 0.17; Fig. 3). As compared to their respective control (formulations without EPN), the efficacy of each EPN formulation was 100 %, 94±18 % or 72±27 % (mean±sd) for gel, SPF or water respectively.

**Figure 3:**
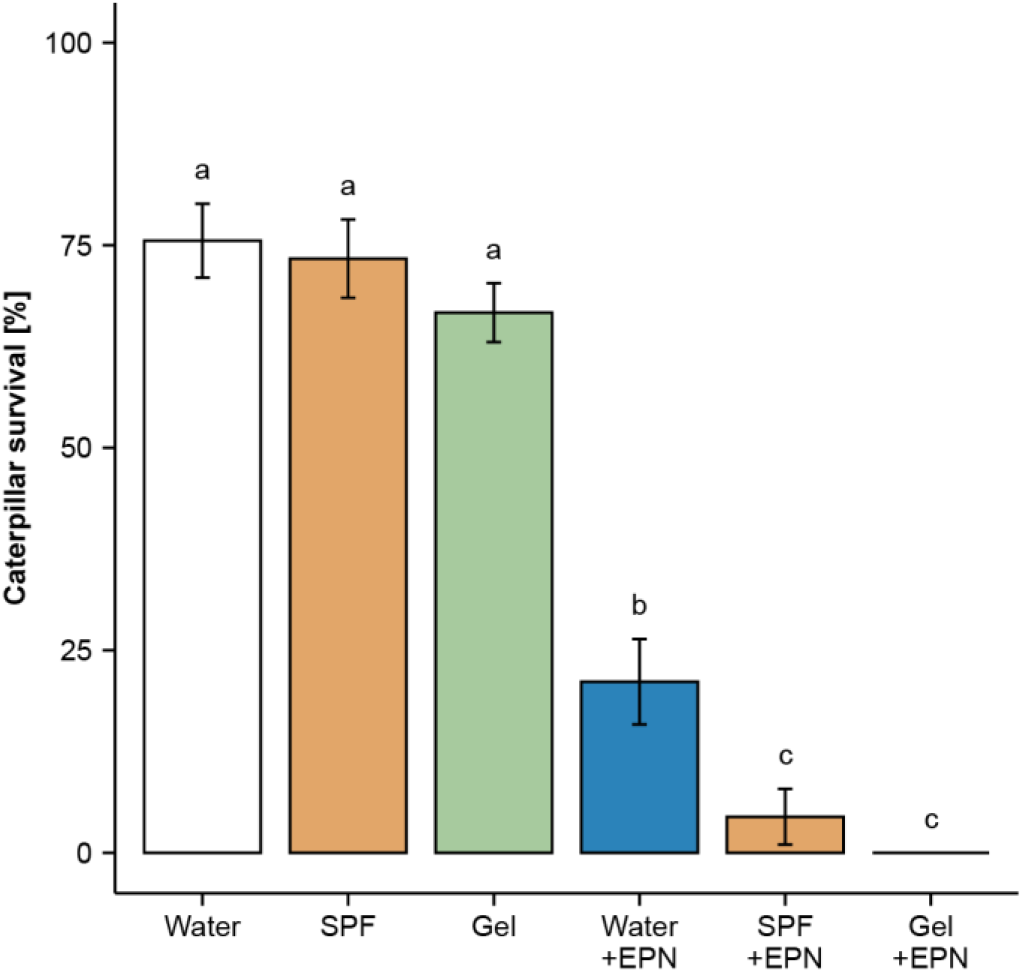
Caterpillar survival (mean±se) six days after applying differently formulated *Steinernema carpocapsae* RW14-G-R3a-2 or formulations without nematodes into the whorl of maize plants. Plants were infested with three third-instar fall armyworms per plant. About 3000 infective juvenile nematodes were applied in 2 ml of formulation per plant. Five cages (each containing two plants) per treatment were used in each of three independent experiment*s* (n = 15 cages; 30 plants per treatment). Letters above bars indicate significant differences (p < 0.05) between treatments according to a pairwise permutation test corrected for false discovery with the Benjamini & Hochberg method (1995).

EPN applied in water, SPF or gel all significantly reduced leaf damage caused by FAW (χ^2^(_5_) = 141, *p* < 0.001; Fig. 4A). These treatments also reduced the proportion of plants with medium to heavy damage (a Davis score higher than three; MaxT = 7.2, *p* < 0.001; Fig. 4B). As compared to their respective control (formulations without EPN), the efficacy of EPN to reduce medium and heavy damage was 100 %, 93% or 39% when applied with SPF, gel or water respectively.

**Figure 4:**
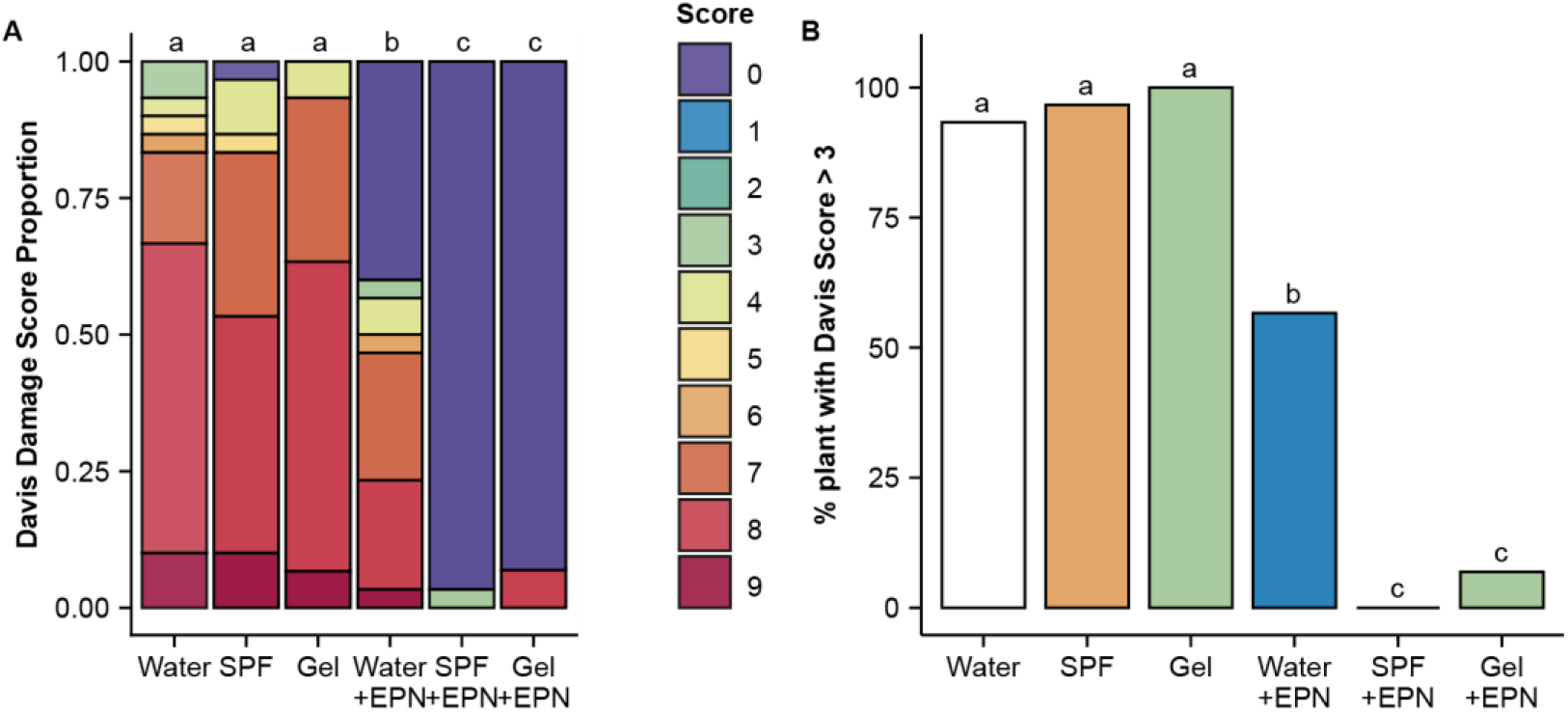
Leaf damage on maize plants six days after applying differently formulated *Steinernema carpocapsae* RW14-G-R3a-2 or formulations without nematodes into leaf whorls. Plants were infested with three third-instar fall armyworms per plant. About 3000 infective juvenile nematodes were applied in 2 ml of formulation per plant. Five cages (each containing two plants) per treatment were used in each of three independent experiments (n = 15 cages; 30 plants per treatment). Plant damage was assessed using the 0 to 9 Davis whorl & furl damage scale, where 0 represent absence of damage and 9 represents a plant almost totally destroyed. (A) Proportion of plants with a given Davis score within treatment. (B) Proportion of plants with medium to heavy damage. Letters above bars indicate significant differences (p < 0.05) between treatments according to (A) multiple comparisons or (B) a pairwise permutation test. Both tests were corrected for false discovery using the Benjamini & Hochberg method (1995).

### 4.2. Efficacy of EPN formulations in reducing fall armyworm infestation and preventing plant damage under field conditions

The treatments affected FAW infestations at both five and ten days post application (five days: χ^2^_(5)_ = 17, *p* = 0.005; ten days: χ^2^_(5)_ = 14, *p* = 0.01; Fig. 5). Five days post treatment, FAW infestation was significantly reduced only by EPN applied in gel (gel+*Sc* vs control: *p* = 0.028; Fig. 5A), and not by any of the other treatments. Ten days post treatment, FAW infestation was significantly reduced by *S. carpocapsae* applied in gel, as well as by cypermethrin (gel+*Sc* vs. control:, *p* = 0.042; cypermethrin: *p* = 0.037; Fig. 5B). As compared to the water control, *S. carpocapsae* applied in gel reduced FAW infestation by 41±15 % [mean±sd] and 34±40 % within five and ten days post treatment, respectively, while cypermethrin achieved 35±29 % and 41±42 % control.

**Figure 5:**
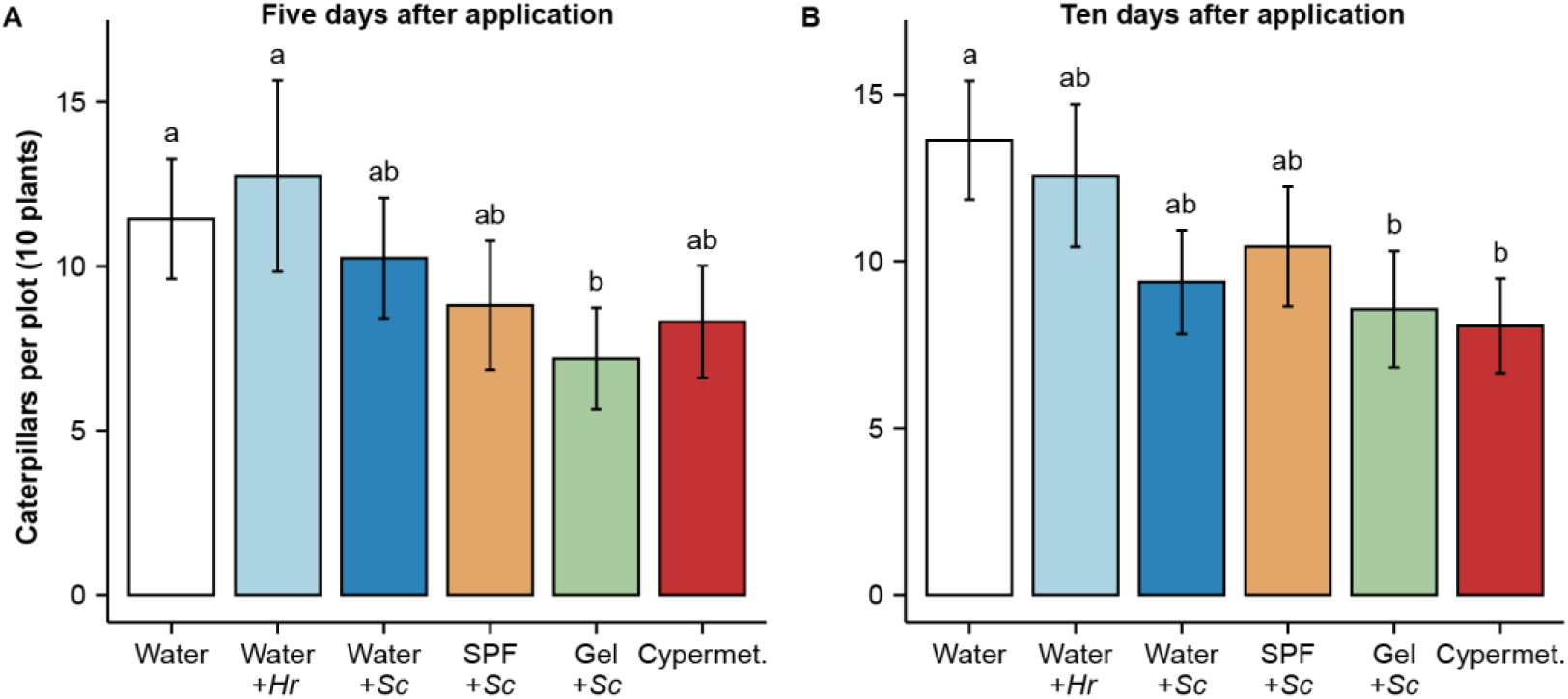
Average number of fall armyworms per plot of ten plans five (A) and ten days (B) after applying differently formulated *Steinernema carpocapsae* RW14-G-R3a-2 (*Sc*) or *Heterorhabditis ruandica* Rw18_M-Hr1a (*Hr*) into leaf whorls of maize, as compared to a commonly used insecticide, cypermethrin (Cypermet.). Four experiments (= maize fields) with natural infestation of fall armyworms (4 plots per treatment per experiment; n = 16 plots per treatment) were carried out in the districts of Nyamagabe and Huye in southern Rwanda in 2020. About 3000 infective juvenile nematodes were applied in 2 ml of formulation per plant. Forty plants were assessed per treatment and per field at both five and ten days post treatment (160 plants per treatment and date). Letters above bars indicate significant differences (*p* < 0.05) according to multiple comparisons corrected for false discovery using the Benjamini & Hochberg method (1995).

Overall, plant damage was significantly affected by the treatments, at both five and ten days post application (five days post treatment: χ^2^_(5)_ = 17, *p* = 0.005; ten days post treatment: χ^2^_(5)_ = 17, *p* = 0.005; Fig. 6A and B). Medium and heavy crop damage were also found to be significantly reduced at both sampling dates (five days post treatment: χ^2^_(5)_ = 12, *p* = 0.04; ten days post treatment: χ^2^_(15)_ = 17, *p* = 0.004; Fig. 6C and D).

**Figure 6:**
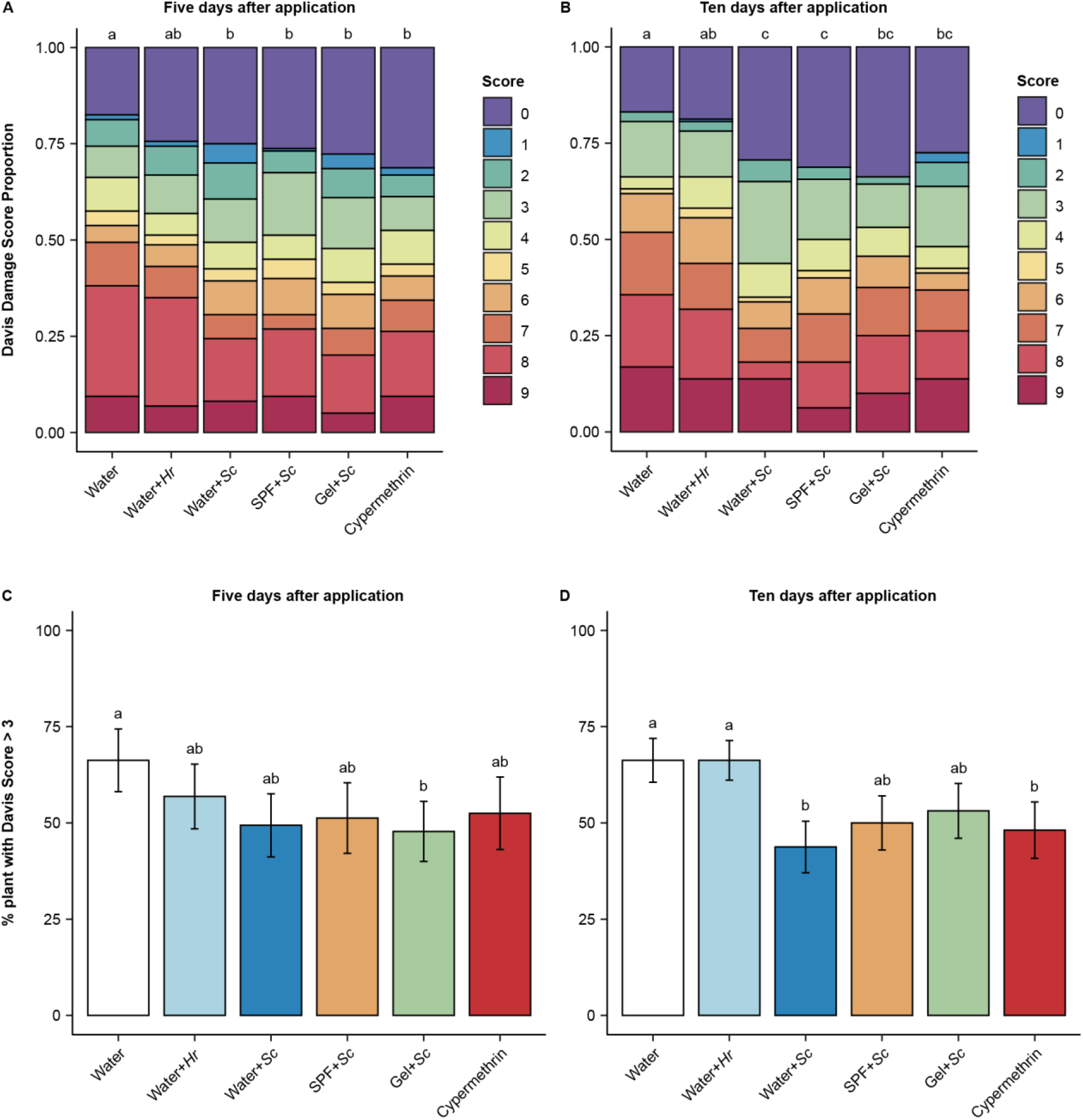
Leaf damage at five (A and C) and ten (B and D) days after applying differently formulated *Steinernema carpocapsae* RW14-G-R3a-2 (*Sc*) or *Heterorhabditis ruandica* Rw18_M-Hr1a (*Hr*) into leaf whorls, as compared to the commonly used insecticide cypermethrin. Four experiments (= maize fields) with natural infestation of fall armyworm caterpillars (4 plots per treatment per experiment, n = 16 plots per treatment) were carried out in the districts of Nyamagabe and Huye in southern Rwanda in 2020. About 3000 infective juvenile nematodes were applied in 2 ml of formulation per plant. Forty plants were assessed per treatment and per field at both five and ten days post treatment (n = 160 plants per treatment and date). (A and B) Proportion of plants within treatments with a given damage score according to the 0 to 9 Davis whorl & furl damage scale, where 0 represent absence of damage and 9 represents a plant that is almost totally destroyed. (C and D) Proportion of plants with medium to heavy damage. Letters above bars indicate significant differences (*p* < 0.05) between treatments according to multiple comparisons corrected for false discovery using the Benjamini & Hochberg method (1995).

Five days post treatment, all the differently formulated *S. carpocapsae* as well as cypermethrin reduced FAW-leaf damage to maize plants (water+*Sc* vs control: *p* = 0.014; SPF+*Sc* vs control: *p* = 0.04; gel+*Sc* vs control: *p* = 0.001; cypermethrin vs control: *p* = 0.014; Fig. 6A). In contrast, *H. ruandica* (water+*Hb* vs control: *p* = 0.19; Fig. 6A) had no effect on leaf damage. *S. carpocapsae* in gel was the only treatment that significantly reduced medium and heavy crop damage (Davis score higher than three: gel+*Sc* vs control: *p* = 0.029; Fig. 6C).

Ten days post treatment, cypermethrin, as well as all formulations with *S. carpocapsae* reduced FAW damage (water+*Sc* vs control*p* = 0.015; SPF+*Sc* vs control: *p* = 0.015; gel+*Sc* vs control: *p* = 0.037; cypermethrin vs control: *p* = 0.043; Fig. 6B), whereas *H. ruandica* had no effect on leaf damage (water+*Hb* vs control: *p* = 0.67; Fig. 6B). Only *S. carpocapsae* applied in water as well as cypermethrin application significantly reduced the proportion of medium and heavily damaged plants (Davis score higher than three: water+*Sc* vs control: *p* = 0.017; cypermethrin vs control: *p* = 0.039; Fig. 6D).

Rapid re-infestation of maize plants by FAW was observed, which was evident from the occurrence of new neonate larvae on the plants. None of the one-time treatments affected these re-infestations (five days post treatment: χ^2^_(5)_ = 5.4, *p* = 0.37; ten days post treatment: χ^2^_(5)_ = 4.5, *p* = 0.48; Fig 7).

**Figure 7:**
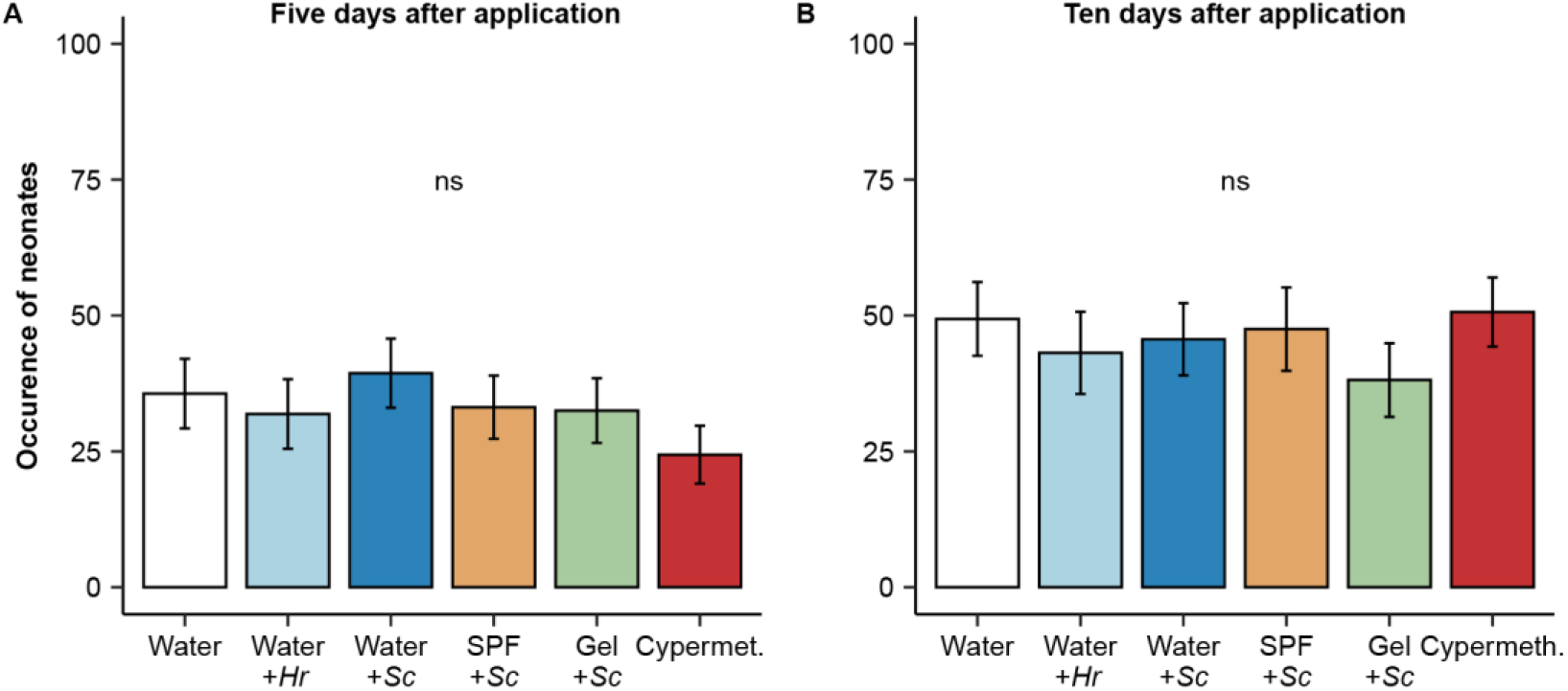
Proportion of plant re-infested with fall armyworm neonates (A) five and (B) ten days after applying differently formulated *Steinernema carpocapsae* RW14-G-R3a-2 (*Sc*) or *Heterorhabditis ruandica* Rw18_M-Hr1a (*Hr*) into leaf whorls, and in comparison to the commonly used insecticide cypermethrin. Four experiments (= maize fields) with natural infestation of fall armyworm caterpillars (4 plots per treatment per experiment, n = 16 plots per treatment) were carried out in the districts of Nyamagabe and Huye in southern Rwanda in 2020. About 3000 infective juvenile nematodes were applied in 2 ml of formulation per plant. Forty plants were assessed per treatment and per field at both five and ten days post treatment (n = 160 plants per treatment and date). No significant differences (ns;*p* > 0.05) were found between treatments using generalized linear mixed-effects models.

## 5. Discussion

EPN have been successfully applied against many pest insects worldwide, and offer a safe alternative solution to synthetic pesticides (Ehlers, 2001; Ehlers and Hokkanen, 1996; Holmes et al., 2015; Kagimu et al., 2017). Here we show that the application of EPN has also great potential for the effective control of FAW caterpillars on maize plants. As FAW larvae are above-ground pests, they rarely encounter soil-borne EPN and are therefore poorly adapted to resist them. This may explain why the caterpillars have been found to be highly susceptible to many species and strains of EPN (Acharya et al., 2020; Fallet et al., in press). EPN normally only reproduce inside an insect host, but they can also be mass produced in fermenters with cultures of their specific symbiotic bacteria or in semi-solid cultures. Hence, these tiny biocontrol agents EPN offer many advantages and it is shown here that the proper application of EPN also holds great promise as a strategy to fight FAW.

In standardised laboratory experimentation, we found that the whorl application of EPN in either a carboxymethyl cellulose gel, a commercial surfactant-polymer-formulation, or just in water killed respectively 100%, 94% or 72% FAW caterpillars on young maize plants (Fig. 3), thereby drastically reducing damage to the plants (Fig. 4). This required a dose of only 3000 IJs per plant. The application of the same formulations without EPN did not affect FAW mortality, showing that the treatment effects were solely due to the EPN. The slightly lower efficacy of EPN applied in water compared to other formulations may be explained by significant run off and relatively rapid evaporation of water and desiccation of the EPN. In contrast, the more viscos gel retains humidity and fills up the whorl, while the commercial adjuvant sticks to the leaf surface, further explaining the higher efficacy of EPN in these two formulations.

The positive results from the laboratory also held true in our field trials. We found the EPN application to be effective against FAW under typical field conditions, even under the high pest infestation levels in the four fields in Rwanda (Fig. 5 and 6). Thirty years ago, Richter and Fuxa (1990) already explored whether EPN can be used against FAW, but obtained inconsistent efficacy under field conditions. They found that the application of *Steinernema feltiae* in water into the whorl of maize seedlings only reduced FAW infestation in one of three experiments (Richter and Fuxa, 1990). Similarly, Garcia et al. (2008) did not observe clear treatment effects when applying *Steinernema* sp. in water into the whorl of artificially infested maize. Using a similar approach, Negrisoli et al. (2010) achieved low efficacy with *S. carpocapsae* and *Heterorhabditis indica*. The addition of tensioactive agents, was not sufficient to improve the efficacy of *Steinernema* sp. (Garcia et al., 2008). We too did not achieve significant control of FAW in field conditions when treating plants with EPN formulated in water. Adding SPF to the EPN helped somewhat (Fig. 5 and 6), but we achieved the best results with EPN in a cellulose gel (Fig. 5, 6 and S1). In fact, *S. carpocapsae* (strain RW14-G-R3a-2) applied in the gel was as effective in killing FAW and reducing leaf damage as the contact pesticide cypermethrin. This pesticide is commonly used against FAW throughout Africa and beyond (Uzayisenga et al., 2020). The specific properties of the here introduced gel formulation appear to contribute to the effectiveness of the EPN in controlling FAW. EPN in just water was the least effective treatment and seeped out of the whorl during application, whereas the gel persisted on the plants for several days.

Not only the EPN-formulation, but also the specific EPN species and strain was found to be important for the successful control of FAW under field conditions. The choice for *S. carpocapsae* (RW14-G-R3a-2) and *H. ruandica* (Rw18_M-Hr1a) for our study was based on extensive laboratory screening in which we compared the virulence of 40 EPN strains, representing twelve species, originating from Mexico, Rwanda, and from a few commercial sources (Fallet et al., in press). Those screenings showed that most EPN species and strains can kill FAW caterpillars, but certain strains were found more infectious than others (Fallet et al., in press). In addition, in the field, the harsh abiotic conditions will affect some species and strains more than others (Hiltpold, 2015; Shapiro-Ilan and Dolinski, 2015), and the differences in EPN killing power may therefore be more variable among strains than in the laboratory.

Although not significant, it appears in the field conditions that *S. carpocapsae* (RW14-G-R3a-2) was more lethal to FAW than *Heterorhabditis ruandica* (Rw18_M-Hr1a) when both species were applied with water. This trend could be explained by the generally high tolerance of *S. carpocapsae* to radiation (Gaugler et al., 1992; Shapiro-Ilan et al., 2015) and desiccation (Brown and Gaugler, 1997; Shapiro-Ilan et al., 2014). EPN behaviour is yet another factor influencing their efficacy in biological control (Hiltpold et al., 2015; Shapiro-Ilan and Dolinski, 2015). Different species and strains of EPN use varying foraging strategies along a continuum from ambushers to crusaders (Selvan et al., 1993). Some EPN, such as *H. ruandica*, are typical crusaders and actively look for their host, whereas others, such as *S. carpocapsae* are ambushers and wait for the host to pass by (Lewis and Clarke, 2012). Considering the crusader strategy of *H. ruandica*, we could have expected that it would actively look for the pest and be more effective. We found the opposite in our field experiments. Possibly, when *H. ruandica* actively searches for hosts on maize plants it is more exposed to lethal abiotic stresses. Evidently, the most appropriate and effective EPN candidate for biological control of the FAW should be carefully selected depending on multiple factors.

The relative high number of FAW caterpillars recovered in the EPN treated plots in our field experiments may have been the result of migration from the untreated buffer plants. Indeed, we observed large numbers of older larvae crawling among plots. This migrating behaviour is common for FAW, especially when plants are heavily infested (Pannuti et al., 2016). With this in mind, our study suggests that just one application of a low dose of *S. carpocapsae* applied in gel can already significantly reduce FAW infestation and prevent heavy damage. However, season-long crop protection may not be possible with just one application, regardless of it being EPN or a synthetic insecticide. Further studies will need to confirm that multiple treatments can indeed fully control FAW. For this, different levels of FAW infestations in different agricultural settings, ranging from small scale African farming to more extensive and commercial farming, should be considered.

In further steps, we also aim to improve the formulation. A first approach would be the incorporation of feeding stimulants to encourage FAW caterpillars to move towards and feed on the EPN-containing substrate. Other additives could protect EPN from harmful abiotic factors such as UV radiation and desiccation. Additional improvements might be achieved by artificial selection of EPN strains for enhanced longevity under field conditions. Selective breeding has been shown to greatly enhance specific traits in EPN, such as tolerance to desiccation and heat and responsiveness to foraging cues (Anbesse et al., 2013; Hiltpold et al., 2010; Mukuka et al., 2010; Perry et al., 2012). Another approach would be to increase the dose of EPN to a level that provides better FAW control. Normally, 2-4 billion EPN are applied per hectare to ensure sufficient pest control (Georgis, 1990; Toepfer et al., 2010), which is in sharp contrast to the ~3000 EPN per plant tested here (representing ca. 0.2-0.3 billion EPN/ha). Our experiments in the laboratory imply that this dose can rapidly achieve 100% mortality of FAW on a plant (Fig. 3). Dose response tests have confirmed the high infectivity of the strains that we used here (Fallet et al., in press). It may therefore be more beneficial to increase the number of applications, and stick to the relative low dose of EPN per application to ensure a low production cost.

In conclusion, our study represents a promising first step towards the development of a safe, sustainable and effective alternative to chemical insecticides. Controlling FAW through the use of formulated EPN seems particularly realistic in an African and Asian context, where low tech manual labour is predominantly used in pest management efforts, allowing manual spot applications in maize fields. Moreover, given the availability of EPN in local soils and the relatively low number of EPN needed to control FAW, we envision that smallholder farmers, provided specific training, could produce their own locally isolated EPN to fight the FAW in a practical, economically viable and environmentally friendly way. The EPN formulations should be adaptable to large-scale high-tech application across commercial maize fields using high wheel precision farming machinery. Regardless of the application technology, we believe our findings clearly underpin the feasibility of using EPN based biocontrol products against FAW.

## Authorship contribution statement

**Patrick Fallet:** Conceptualization, Methodology, Formal analysis, Investigation, Data Curation, Writing - Original Draft, Visualization, Supervision, Project administration. **Didace Bazagwira:** Investigation, Writing - Review & Editing. **Julie Guenat:** Investigation, Data Curation, Writing - Review & Editing. **Carlos Bustos Segura:** Methodology, Formal analysis, Writing - Review & Editing. **Patrick Karangwa:** Resources, Writing - Review & Editing, Project administration. **Ishimwe Primitive Mukundwa:** Investigation, Writing - Review & Editing. **Joelle Kajuga:** Investigation, Resources, Writing - Review & Editing, Project administration. **Thomas Degen:** Visualization, Writing - Review & Editing. **Stefan Toepfer:** Conceptualization, Methodology, Investigation, Supervision, Writing - Review & Editing, Project administration, Funding acquisition. **Ted C. J. Turlings:** Conceptualization, Methodology, Supervision, Writing - Review & Editing, Project administration, Funding acquisition.

## De claration of Competing Interest

The authors declare that they have no known competing financial interests or personal relationships that could have appeared to influence the work reported in this paper.

## Acknowledgment

The authors would like to thank the Rwanda Agriculture and Animal Resource Development Board for collaborative nematodes surveys and research, including benefit sharing, material transfer agreements, and research permits. The authors are grateful to Patrick Grof-Tisza and Radu Alexandru Slobodeanu for their advice on statistical analysis, Arnaud Feutrier for his help in the field experiments, Anant V. Patel for his suggestions regarding the gel formulations. The authors thank Ralf-Udo Ehlers and Bart Vandenbossche for kindly providing us with the commercial surfactant-polymer-formulation.

## Supplementary material

**Figure S1:**
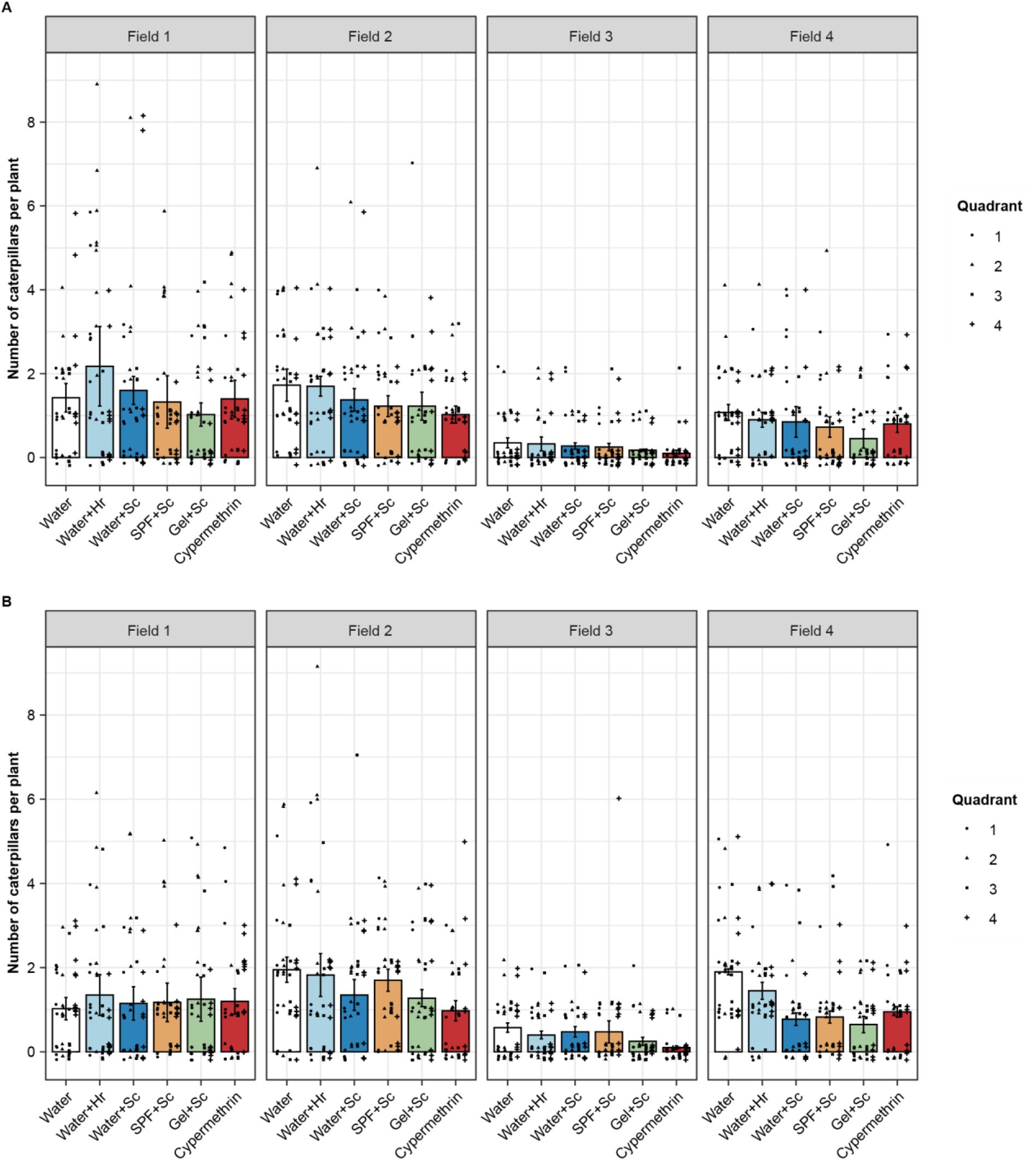
Number of fall armyworm caterpillars per plant within each field, five (A) and ten days (B) after applying differently formulated *Steinernema carpocapsae* RW14-G-R3a-2 (*Sc*) or *Heterorhabditis ruandica* Rw18_M-Hr1a (*Hr*) into leaf whorls of maize, as compared to a commonly used insecticide, cypermethrin (Cypermethtin). Bars represent the average number of caterpillars per plant within each field and treatment. Dots represent the number of caterpillars per plant within each field, treatment and quadrant. Four experiments (= maize fields) with natural infestation of fall armyworms caterpillars (4 plots per treatment per experiment). Treatment consisted of about 3000 infective juvenile nematodes applied in 2 ml of formulation per plant. Forty plants were assessed per treatment and per field at both five and ten days post treatment (n = 160 plants per treatment and date).

**Figure S2:**
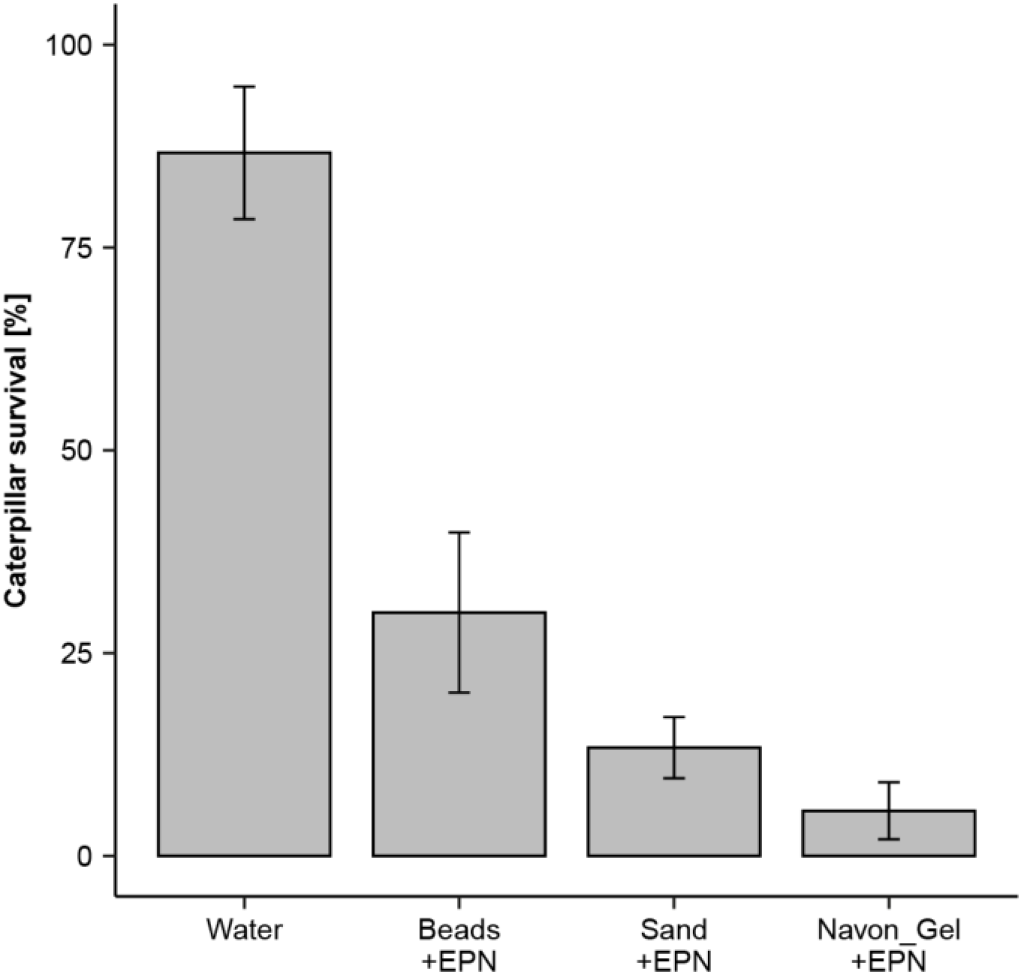
Caterpillar survival (mean±se) six days after applying differently formulated *Steinernema carpocapsae* RW14-G-R3a-2 into the whorl of maize plants. Plants were infested with three third-instar fall armyworms per plant. About 3000 infective juvenile nematodes were applied in 1 ml of formulation per plant. Five cages (each containing two plants) per treatment were used in one experiment (n = 5 cages; 10 plants per treatment).

**Figure S3:**
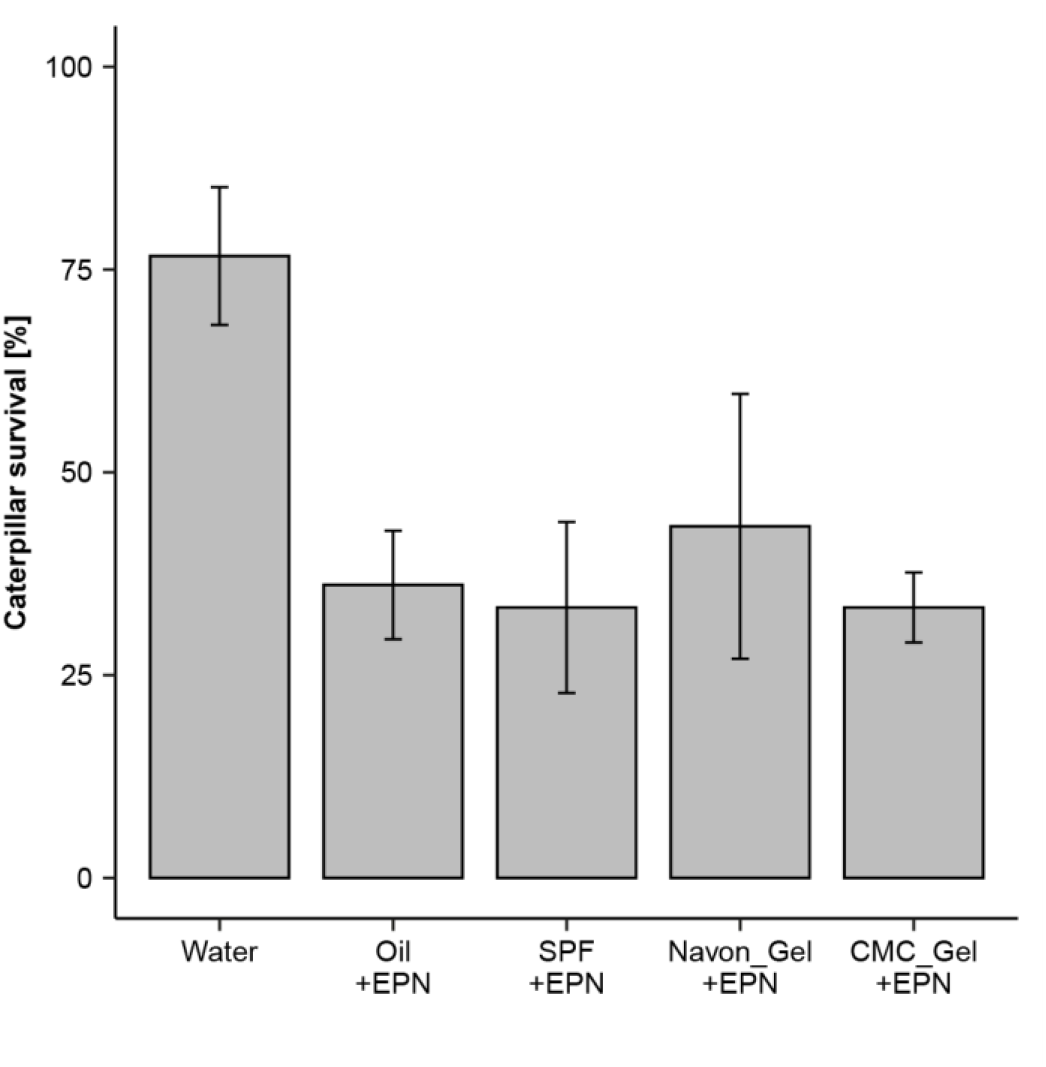
Caterpillar survival (mean±se) six days after applying differently formulated *Heterorhabditis ruandica* Rw18_M-Hr1a into the whorl of maize plants. Plants were infested with three third-instar fall armyworms per plant. About 3000 infective juvenile nematodes were applied in 1 ml of formulation per plant. Six cages (each containing two plants) per treatment were used in one experiment (n = 6 cages; 12 plants per treatment).

**Figure S4:**
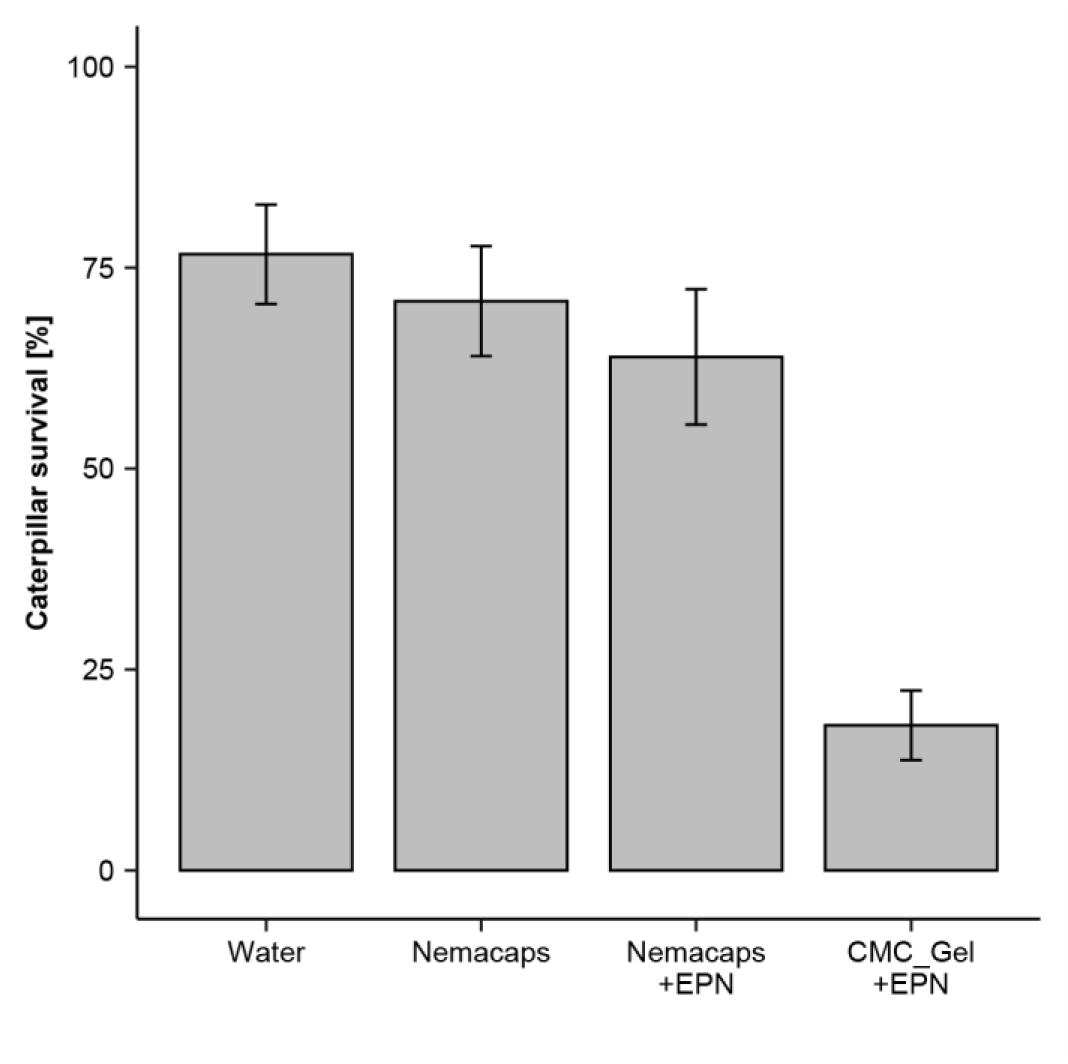
Caterpillar survival (mean±se) six days after applying differently formulated *Heterorhabditis ruandica* Rw18_M-Hr1a into the whorl of maize plants. Plants were infested with three third-instar fall armyworms per plant. About 3000 infective juvenile nematodes were applied in 1 ml of formulation per plant. Six cages (each containing two plants) per treatment were used in each of two independent experiments (n = 12 cages; 24 plants per treatment).

**Figure S5:**
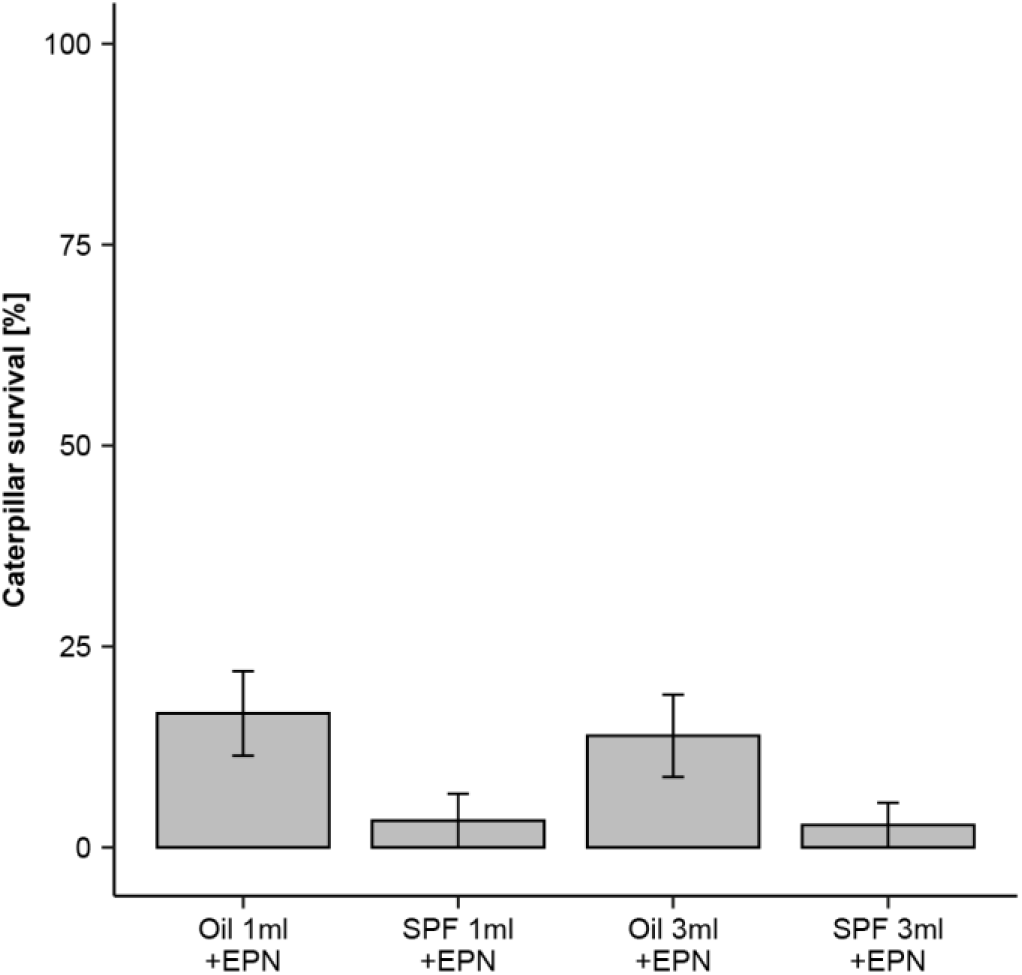
Caterpillar survival (mean±se) six days after applying differently formulated *Heterorhabditis ruandica* Rw18_M-Hr1a into the whorl of maize plants. Plants were infested with three third-instar fall armyworms per plant. About 3000 infective juvenile nematodes were applied in 1 or 3 ml of formulation per plant. Five cages (each containing two plants) per treatment were used in one experiment (n = 5 cages; 10 plants per treatment).

## Equations

With the laboratory data, the efficacy of each formulation in killing FAW was estimated as follows:

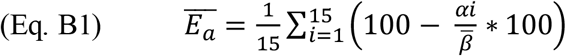

where, αi is the number alive caterpillars in the treated cage i, and

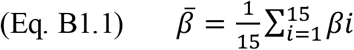

is the average number of alive caterpillars in the control cages, with βi is the number of caterpillars in the control cage i.

With the laboratory data, the efficacy of each formulation in preventing medium to heavy damage was estimated as follows:

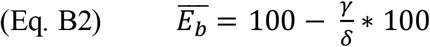

where, γ is the number of damaged plants in the treatment, and δ is the number of damaged plants in the control.

With the field data, the efficacy of each treatment in reducing FAW infestation was estimated as follows:

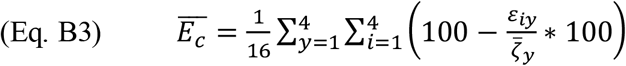

where, ε_iy_ is the number of caterpillars in the treated plot i in field y, and

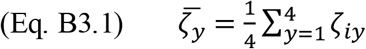

is the average number of caterpillars in the control plots of field y, with ζ_iy_ is the number of caterpillars in the control plot i in field y.

